# A happy accident: a novel turfgrass reference genome

**DOI:** 10.1101/2022.03.08.483531

**Authors:** Alyssa R. Phillips, Arun S. Seetharam, Patrice S. Albert, Taylor AuBuchon-Elder, James A. Birchler, Edward S. Buckler, Lynn J. Gillespie, Matthew B. Hufford, Victor Llaca, M. Cinta Romay, Robert J. Soreng, Elizabeth A. Kellogg, Jeffrey Ross-Ibarra

## Abstract

*Poa pratensis*, commonly known as Kentucky bluegrass, is a popular cool-season grass species used as turf in lawns and recreation areas globally. Despite its substantial economic value, a reference genome had not previously been assembled due to the genome’s relatively large size and biological complexity that includes apomixis, polyploidy, and interspecific hybridization. We report here a fortuitous *de novo* assembly and annotation of a *P. pratensis* genome. Instead of sequencing the genome of a C4 grass, we accidentally sampled and sequenced tissue from a weedy *P. pratensis* whose stolon was intertwined with that of the C4 grass. The draft assembly consists of 6.09 Gbp with an N50 scaffold length of 65.1 Mbp, and a total of 118 scaffolds, generated using PacBio long reads and Bionano optical map technology. We annotated 256K gene models and found 58% of the genome to be composed of transposable elements. To demonstrate the applicability of the reference genome, we evaluated population structure and estimated genetic diversity in *P. pratensis* collected from three North American prairies, two in Manitoba, Canada and one in Colorado, USA. Our results support previous studies that found high genetic diversity and population structure within the species. The reference genome and annotation will be an important resource for turfgrass breeding and study of bluegrasses.

## Introduction

*Poa pratensis* L., commonly known as Kentucky bluegrass, is an economically valuable horticultural crop grown globally on lawns and recreational areas as turf (Haydu *et al*. 2006). Native to Europe and Asia, it was introduced to North America in the seventeenth century by European colonizers as a forage crop (Carrier and Bort 1916; Raggi *et al*. 2015). Today, Kentucky bluegrass is the most popular cool-season grass used for turf due to it’s vigorous growth and quick establishment that creates a dense, strong sod with a long lifespan (Casler and Duncan 2003).

Today, there are 40 million acres of managed turf in the United States (U.S.), an area approximately the size of the state of Florida (Milesi *et al*. 2005). While this massive area has the potential to serve as an important carbon sink, the large water and fertilization resources required currently outweigh the benefits (Milesi *et al*. 2005). Breeding efforts are underway to improve environmental-stress tolerances, disease and insect resistance, seed quality and yield, as well as uniformity and stability of traits (reviewed in Bonos and Huff 2013). While the economic value of *P. pratensis* is high, it is highly invasive, and in the last 30 years has aggressively invaded the North American Northern Great Plains, altering ecosystem function by reducing pollinator and plant diversity and altering nutrient dynamics (Kral-O’Brien *et al*. 2019; DeKeyser *et al*. 2015; Hendrickson *et al*. 2021). Continued research into the genetic diversity of wild *P. pratensis* is needed to understand how invasive populations are rapidly adapting, and the study of wild populations may enable identification of disease or environmentally tolerant ecotypes for use in turfgrass breeding.

Previous studies using RAPD, ISSR, and SRR markers demonstrated high genetic diversity in both developed cultivars and wild populations but limited population structure between these groups (Bushman *et al*. 2013; Raggi *et al*. 2015; Honig *et al*. 2012, 2018, but see Dennhardt *et al*. 2016). Population divergence has been detected amongst some wild populations (Dennhardt *et al*. 2016) but the extent of population structure is unclear. There are a number of potential reasons for finding a lack of population structure, including gene flow, the independent development of cultivated lines from locally adapted ecotypes (Raggi *et al*. 2015; Bonos and Huff 2013), and geographic heterogeneity in patterns of genetic diversity. Repeated reversion of cultivars to wild forms has also been suggested, but is unlikely (Dennhardt *et al*. 2016). Alternatively, previous studies may simply not have had sufficient marker resolution to detect population structure in a highly heterozygous polyploid species like *P. pratensis*.

Genetic analysis and improvement of turfgrass are challenging because of apomixis and polyploidy (Bushman and Warnke 2013). *Poa pratensis* is a facultative apomict, meaning it can reproduce sexually or asexually by aposporous apomixis, and it is a polyploid with frequent aneuploidy (Brown 1939). Although apomixis is a highly valued trait for seed production, high rates of apomixis stymie the recombination needed to genetically analyze traits or recombine beneficial traits into one cultivar (Bonos and Huff 2013). Polyploidy and aneuploidy further these difficulties due to copy number variation of regions of interest and non-Mendelian inheritance resulting from double reduction. While some progress has been made in managing apomixis (Funk *et al*. 1967; Pepin and Funk 1971; Matzk 1991), including the discovery of its genetic basis (Albertini *et al*. 2004; Marconi *et al*. 2020), the development of additional molecular and genomic tools in *P. pratensis* are needed to move genetic analysis and breeding efforts forward in the face of its complex biology.

Here, we report the first *P. pratensis* genome. While attempting to assemble the genome for a C4 prairie grass, *Andropogon gerardi*, we unknowingly sequenced and assembled a wild *Poa* growing in the same pot. Fortunately, this resulted in a highly contiguous, near complete genome assembly. We utilized the reference genome and wild *Poa* from three prairies to investigate the genetic diversity and population structure of North American *Poa*. The reference genome and annotation presented here are an important advancement for Kentucky bluegrass breeding. Additionally, this reference genome provides an important resource for the study of closely related bluegrasses including *P. trivialis* L., *P. annua* L., and *P. arachnifera* Torr.

## Materials and Methods

### Sample collection

Rhizomes of *Poa* species were collected fortuitously as part of a different project aimed at collecting major C4 prairie grasses (*Andropogon gerardi* Vitman, *Sorghastrum nutans* (L.) Nash, and *Schizachyrium scoparium* (Michx.) Nash) in moist prairies in Colorado, USA and two prairies in Manitoba, Canada and (Supplement 1). Necessary permissions and permits were obtained prior to collecting. Plants were brought back to the United States from Canada under phytosanitary certificate 3193417.

The C4 focal plants were dug up with a shovel late in the growing season in 2018 (when the *Poa* was dormant and thus invisible), soil was washed off, rhizomes were wrapped in wet paper towels, and leaves were cut back to about 4 inches height to reduce transpiration. The focal C4 plant was placed in a 1gallon Ziploc bag and returned to the plant growth facility at the Donald Danforth Plant Science Center in St. Louis, MO, USA. Plants were potted in 2:1 BRK20 promix soil to turface. The previously dormant *Poa* plants produced fresh green leaves in this setting and grew faster than the C4 plant with which it was entwined. Once it was discovered that *Poa* had interpolated itself into the rhizome and root area of the C4 plants, the *Poa* plants were extricated and placed in separate pots.

One *Poa* was found inside the pot for an *Andropogon gerardi* genotype which was used to attempt assembly of a reference genome. Instead of collecting tissue from the *A. gerardi* plant, tissue was accidentally sampled from the *Poa* plant. This *Poa* individual is referred to as the *Poa* reference individual (Supplement 1). Eight additional *Poa*, referred to here as the *Poa* population panel, were discovered in various pots for C4 grasses whose genomes we attempted to sequence.

As *Poa* species generally require vernalization to flower, several plants were over-wintered outside under mulch and flowered in spring 2020 and/or 2021; voucher specimens were taken from these plants to verify species identity and have been deposited at the Smithsonian Institution (Washington, District of Columbia, U.S.A) and the Missouri Botanical Garden (St. Louis, MO, U.S.A.) (Heide 1994). Not all *Poa* individuals survived, so some specimens lack vouchers. Additionally, not all surviving *Poa* flowered so vegetative vouchers were submitted (Supplement 1).

### PacBio sequencing

Approximately 4.1 g fresh tissue from the reference individual was extracted for PacBio sequencing using a High Molecular Weight (HMW) DNA approach based on the Circulomics Big DNA Kit (Circulomics, USA). This method yields DNA with a center of mass at 200 Kb, which is sufficient to construct PacBio CLR 20 Kb+ libraries. Sequencing was completed on the Sequel II across four SMRTCells. DNA extraction and sequencing was completed by Corteva Agriscience™.

### Bionano optical map generation

DNA was extracted from 0.7 g of fresh leaf tissue from the reference individual using agarose embedded nuclei and the Bionano Prep™ Plant Tissue DNA Isolation kit. DNA extraction, labeling, imaging, and optical map assembly followed the methods previously described in Hufford *et al*. (2021) and was completed by Corteva Agriscience™.

### Preparation and imaging of metaphase spreads

Metaphase spreads were utilized to estimate chromosome count and ploidy of the reference individual. Root tips were harvested from a recent off-shoot of the reference individual, treated with nitrous oxide (3 hr at 160 psi) to stop mitosis in metaphase (Kato 1999), then processed as previously described in Kato *et al*. (2004) and Kato *et al*. (2011) with minor modification. Specifically, the root tips were fixed in 90% acetic acid for 15 min, then rinsed with and stored in 70% ethanol at − 20^*◦*^C. Ethanol was removed from the root tips prior to enzymatic digestion by soaking in water for 10 min. About 1 mm of the tip (meristem and root cap) was excised and transferred to a tube containing 20 µL of 3% cellulase R-10 (Desert Biologicals, Phoenix, AZ) and 1.25% pectolyase Y-23 (Desert Biologicals) in citrate buffer (10 mM sodium citrate, 10 mM EDTA, adjusted to pH 5.5 with citric acid) on ice. The tissue was digested for approximately 1 hr at 37^*◦*^C. Seventy percent ethanol was used to inactivate the enzymes and rinse the samples. The ethanol was replaced with approximately 7 µL of a solution of 90% acetic acid and 10% methanol. The tissue was broken and cells dispersed using a blunted dissecting probe. The entire volume was dropped from a height less than 1 cm onto a microscope slide in a container lined with wet paper towels and allowed to dry.

Preparations were counterstained with a 1/20 dilution of Vectashield with DAPI (Vector Laboratories, Burlingame, CA). Images were captured using Applied Spectral Imaging software (Carlsbad, CA) on an Olympus BX 61 fluorescence microscope. Photoshop Brightness/Contrast and Curves functions were used to decrease background noise and better define the chromosomal arms.

### Genome size estimation

Genome size was estimated for the *Poa* reference individual and 4 of the population panel individuals (Supplement 5). Not all population panel individuals were sampled as some plants died prior to estimation. Genome size estimation methods using an internal standard are modified from Doležel *et al*. (2007). Two internal standards were used for the reference: maize B73 inbred line (5.16 pg/2C) and *Andropogon gerardi* accession CAM 1351 (6.13 pg/2C). Only the maize B73 internal standard was used for the population panel. Approximately 10×1 cm of fresh leaf tissue for the target and sample standard were placed in a plastic square petri dish. A chopping solution composed of 1 mL LB01 buffer solution, 250µL PI stock (2 mg/mL), and 25 µL RNase (1 mg/mL) added to the dish (1.25 mL; Doležel *et al*. 2007). The tissue was then chopped into 2-4 mm lengths and the chopping solution was mixed through the leaves by pipetting. The solution was then pipetted through a 30µm sterile single-pack CellTrics® filter into a 2 mL Rohren tube on ice. Three replicates were chopped separately and analyzed for each *Poa* population panel genotype and 9 replicates were analyzed for the reference. The samples were left to chill for 20 min before analysis with a BD Accuri™ C6 flow cytometer. Samples were run in Auto Collect mode with a 5-min run limit, slow fluidics option, a FSC-H threshold with less than 200,000 events, and a 1-cycle wash. The cell count, coefficient of variation of FL2A, and mean FL2-A were recorded for the target and reference sample with no gating. Results were analyzed separately for each replicate and manually annotated to designate the set of events. The replicates for each *Poa* genotype were averaged (Supplement 6).

### Illumina sequencing of the Poa population panel

DNA was extracted from the *Poa* population panel using approximately 100 mg of lyophilized leaf tissue and a DNeasy® Plant Kit (Qiagen Inc., Germantown, MD). High throughput Illumina Nextera ® libraries were constructed and samples were sequenced with other plant samples in pools of 96 individuals in one lane of an S4 flowcell in an Illumina Novaseq 6000 System with paired-end 150-bp reads, providing approximately 0.80X coverage for each sample.

### Species identification

Species identification was completed using both morphological and DNA sequence data. Morphological assessment was completed for the *Poa* reference genome and three of the population panel samples using flowering and vegetative vouchers. Phylogenetic inference was completed for species identification of all samples using one plastid and two nuclear ribosomal DNA loci: *trn*T-*trn*L-*trn*F (TLF), external transcribed spacer (ETS), and internal transcribed spacer (ITS), respectively. Trees for *mat*K and *rpo*B-*trn*C were also evaluated but the sequences showed little variation across sampled species.

Sequences for these loci were extracted from the *Poa* population panel whole genome sequence data by aligning reads to a *P. pratensis* sequence for each locus downloaded from Genbank (Supplement 2) using the default options of bwa mem (v0.7.17; Li 2013). The alignment files were sorted using SAMtools (v1.7; Danecek *et al*. 2021), read groups were added using Picard AddOrReplaceReadGroups, and duplicates removed with Picard MarkDuplicates using default settings (http://broadinstitute.github.io/picard). We identified variable sites for each sample separately using GATK (v4.1) HaplotypeCaller with default options (Van der Auwera and O’Connor 2020). SNPs were filtered to remove sites with low mapping quality and low sequencing quality (`gatk VariantFiltration-filter “QUAL < 40.0”-filter “MQ < 40.0”` and default gatk SelectVariants). A consensus sequence for each locus and sample was generated using GATK FastaAlternateReferenceMaker, which replaces the gene reference bases at variable sites with the alternate allele.

Sequences were extracted from the reference genome by aligning the *P. pratensis* reference sequences downloaded from Genbank to the reference genome with bwa mem using default options (v0.7.17; Li 2013). This allowed us to identify the position of each locus in the reference. Each locus only mapped to a single region in the reference genome, which was extracted using bioawk (https://github.com/lh3/bioawk).

Sequences from the reference genome and the population panel were included in a dataset with 119 *Poa* samples from previous work (Supplement 3; Cabi *et al*. 2016, 2017; Gillespie *et al*. 2007, 2008, 2009, 2018; Giussani *et al*. 2016; Refulio-Rodriguez *et al*. 2012; Soreng and Gillespie 2018; Soreng *et al*. 2015, 2017, 2020; Sylvester *et al*. 2021). These samples were chosen to represent the phylogenetic diversity of the genus *Poa*, and include all seven currently recognized subgenera as well as 29 of 38 sections and several unclassified species groups (classification according to Gillespie *et al*. (2007), with updates by Cabi *et al*. (2017); Gillespie *et al*. (2008, 2018); Soreng and Gillespie (2018); Soreng *et al*. (2020)). Since formal infrageneric taxonomic delimitations are often imperfect, and the genus *Poa* is large and highly complex, genotype codes are used in Supplement 3 as shorthand for the plastid and nrDNA clades found in a sample or species (see Soreng *et al*. (2020) for the most recent iterations).

Sequences were aligned using the auto-select algorithm and default parameters in the MAFFT plugin (v7.017; Katoh and Standley 2013) in Geneious (v8.1.9; http://www.geneious.com) followed by manual adjustment. *Poa* sect. *Sylvestres* was used as the outgroup to root trees based on its strongly supported position as sister to all other *Poa* species in previous plastid analyses (Gillespie *et al*. 2007, 2009, 2018). Bayesian Markov chain Monte Carlo analyses were conducted in MrBayes (v3.2.6; Ronquist *et al*. 2012). Optimal models of molecular evolution were determined using the Akaike Information Criterion (AIC; Akaike 1974) conducted through likelihood searches in jModeltest (Darriba *et al*. 2012) with default settings. Models were set at GTR + Γ for ETS and GTR + I + Γ for ITS and TLF based on the AIC scores and the models allowed in MrBayes. Two independent runs of four chained searches were performed for three or four million generations, sampling every 500 generations, with default parameters. Analyses were stopped when an average standard deviation of split frequencies of 0.007001, 0.006350, and 0.006490 was reached for ITS, ETS, and TLF, respectively. A 25% burn-in was implemented prior to summarizing a 50% majority rule consensus tree and calculating Bayesian posterior probabilities. Trees were visualized and annotated in R using ggtree (v2.0.4) with ape (v5.4) and treeio (v1.10) (Yu 2020; R Core Team 2017; Wang *et al*. 2020; Paradis and Schliep 2019).

### Genome assembly

PacBio subreads obtained as BAM files were converted to FASTA format using SAMtools (v1.10; Danecek *et al*. 2021) and errorcorrection was performed using overlap detection and error correction module (first stage) of Falcon (v1.8.0; Chin *et al*. 2016). For running Falcon, the following options were used: the expected genome size was set to 6.4 Gbp (-genome_size = 6400000000), a minimum of two reads, maximum of 200 reads, and minimum identity of 70% for error corrections (–min_cov 2 –max_n_read 200, –min_idt 0.70), using the 40x seed coverage for autocalculated cutoff. The average read correction rate was set to 75% (-e 0.75) with local alignments at a minimum of 3000 bp (-l 3000) as suggested by the Falcon manual. For the DAligner step, the exact matching length of k-mers between two reads was set to 18 bp (-k 18) with a read correction rate of 80% (-e 0.80) and local alignments of at least 1000 bp (-l 1000). Genome assembly was performed with Canu (v1.9: Koren *et al*. 2017) using the error-corrected reads from Falcon. For sequence assembly, the corrected reads had over 70x coverage for the expected genome size of *Poa* and were characterized by N50 of 25.6 Kbp and average length of 16.3 Kbp. These reads were trimmed and assembled with Canu using the default options except for ovlMerThreshold=500.

The Canu generated contig assembly was further scaffolded utilizing the Bionano optical map with Bionano Solve (v3.4) and Bionano Access (v1.3.0), as described previously by Hufford *et al*. 2021. The default config file (hybridScaffold_DLE1_config.xml) and the default parameters file (optArguments_nonhaplotype_noES_noCut_DLE1_saphyr.xml) were used for the hybrid assembly. The scaffolding step of Bionano Solve incorporates three types of gaps: 1) gaps of estimated size (varying N-size, but not 100bp or 13bp), using calibrated distance conversion of optical map to basepair (cases when contiguous optical map connects two contigs); 2) gaps of unknown sizes (100-N gaps), when distance could not be estimated (cases when large repeat regions like rDNA or centromeres interrupt the optical map but evidence to connect the map is present); and 3) 13-N gaps, in regions where two or more independently assembled contigs align to the same optical map, overlapping at the ends. The 13-N gaps are usually caused by sequence similarity sufficient for aligning to the optical map, but less than required to merge contigs. This could be caused by either high heterozygosity in that region, highly repetitive sequence, paralogous regions of the sub-genomes, or assembly errors. The contig overlaps, regardless of the size, are connected end-to-end by adding 13-N gaps when processed using Bionano Solve. Due to the polyploid nature of Poa as well as its high heterozygosity, these 13-N gaps had to be manually curated. We inspected the contig alignments to the optical map using using Bionano Access (v1.3.0), either to trim the overlapping sequence or to remove exact duplicates to generate error-free assembly.

### Genome annotation

Gene prediction was carried out using a comprehensive method combining *ab initio* predictions (from BRAKER v2.1.6; Brůna *et al*. 2021) with direct evidence (inferred from transcript assemblies) using the BIND strategy (Li *et al*. 2021). Briefly, 58 RNA-seq libraries were downloaded from NCBI (Supplement 4) and mapped to the genome using a STAR (v2.5.3a; Dobin *et al*. 2013)-indexed genome and an iterative two-pass approach under default options to generate mapped BAM files. BAM files were used as input for multiple transcript assembly programs to assemble transcripts: Class2 (v2.1.7; Song *et al*. 2016), Cufflinks (v2.2.1; Trapnell *et al*. 2012), Stringtie (v2.1.4; Pertea *et al*. 2015) and Strawberry (v1.1.2; Liu and Dickerson 2017). Redundant assemblies were collapsed and the best transcript for each locus was picked using Mikado (v2.3.3; Venturini *et al*. 2018) by filling in the missing portions of the ORF using TransDecoder (v5.5.0; Haas *et al*. 2013) and homology as informed by the NCBI BLASTX (v2.10.1+; Altschul *et al*. 1990) results to the SwissProtDB (Duvaud *et al*. 2021). Splice junctions were also refined using Portcullis (v1.2.1; Mapleson *et al*. 2018) to identify isoforms and to correct misassembled transcripts. Both *ab initio* and direct evidence predictions were analyzed with TESorter (v1.3.0; Zhang *et al*. 2019) to identify and remove any TE-containing genes before merging them. Merging was done using the GeMoMa (v1.8) Annotation Filter tool, to combine and filter gene predictions from BRAKER, Mikado and additional homology-based gene predictions generated by the GeMoMa pipeline using *Hordeum vulgare* annotations (Mascher *et al*. 2021; Keilwagen *et al*. 2016, 2018). The predictions were prioritized using weights, with highest for homology (1.0), followed by direct evidence (0.9) and lowest for gene predictions from *ab initio* methods (0.1). Homology is defined by GeMoMa as protein sequence similarity and and intron position conservation relative to *Hordeum vulgare*. The Annotation Filter tool was run with settings to enforce the completeness of the prediction (start==’M’ stop==’*’), external evidence support (score/aa>=0.75), and RNAseq support (evidence>1 or tpc==1.0). The final predictions were subjected to phylostratiography analyses using phylostratr (v0.20; Arendsee *et al*. 2019). The focal species were set as ‘4545’ for *Poa pratensis*, and default options were used. The program creates a clade tree of species based on the current NCBI tree of life, trims the tree to maximize evolutionary diversity, retrieves the species proteome from Uniprot, and compares the proteins of the focal species to those of other species in the tree using pairwise BLASTs (Diamond search). Each gene is then assigned to the deepest clade in which it has an inferred homolog. Genes found only in the focal species are considered orphan genes and assigned to the phylostratum ‘*Poa pratensis*.’ Final gene-level annotations were saved in GFF3 format and the predicted peptides/CDS sequences were extracted using gffread of the Cufflinks package (v2.2.1; Trapnell *et al*. 2012).

### Assessment of the assembly

Genome contiguity statistics were computed using the Assemblathon script (Bradnam *et al*. 2013). Gene space completeness was measured using BUSCO (v4.0; Manni *et al*. 2021) using the liliopsida_odb10 profile (n = 3278) and poales_odb10 profile (n = 4896) with default options. The contiguity of TE assembly was then assessed using the LTR Assembly Index (LAI; Ou *et al*. 2018). To compute LAI, we first annotated repeats using the Extensive *de-novo* TE Annotator (EDTA; v1.9.6; Ou *et al*. 2019), and intact LTR retrotransposons (LTR-RT) were identified using LTRharvest (v1.6.1; Manchanda *et al*. 2020), and LTR_FINDER_parallel (v1.1; Ellinghaus *et al*. 2008). LTR_retriever (v2.9.0; Ou *et al*. 2018) was then used to filter the intact LTRs and computed the LAI score for the genome.

### Population genetics of Poa

The population panel was mapped to the scaffold assembly, excluding the alternate scaffolds, using bwa mem (v0.7.17; Li 2013). Reads were sorted using SAMtools (v1.7; Danecek *et al*. 2021), read groups were added using Picard AddOrReplaceReadGroups, and duplicates removed with Picard MarkDuplicates (http://broadinstitute.github.io/picard) using default settings.

Site filtering and genotyping was completed with ANGSD (v0.934; Korneliussen *et al*. 2014). Reads were filtered, retaining unique reads, reads with a flag below 255, and proper pairs (angsd-uniqueOnly 1 -remove_bads 1 -only_proper_pairs 1 -trim 0), as well as a minimum mapping and base quality of 30 (angsd -minMapQ 30 -minQ 30). Sites were filtered with a strict maximum depth cutoff in order to exclude sites where paralogs may be mapping. Assuming read depth follows a Poisson distribution with a mean of 0.8, we expect 99% of reads to have a depth of 4 or less. We included sites with a minimum depth of 1 and a maximum depth of 4 and required all genotypes to have data at a site (angsd –doCounts 1 -setMinDepthInd 1 -setMaxDepthInd 4 -minInd 8). Sites were also filtered for a minor allele frequency greater than 5% in the principal component analysis (PCA; angsd –doMajorMinor 4 -doCounts 1 -doMaf 1 -minMaf 0.05).

After filtering, a single-read was randomly sampled at each base to serve as the genotype (angsd -doIBS 1). This genotyping approach is discussed in Results and Discussion. A genotype matrix was sampled three independent times for each of the following analyses in order to assess sampling error.

Population structure and nucleotide diversity were evaluated to demonstrate the utility of the *P. pratensis* reference genome. Population structure was assessed using a principal component analysis (PCA) implemented in ANGSD (angsd -doCov). A PCA was run with all *Poa* and only *P. pratensis*. The covariance matrices were plotted with ggplot2 (v3.4) in R (R Core Team 2017; Wickham 2016)

Nucleotide diversity was estimated for each *P. pratensis* genotype in the *Poa* population panel as nucleotide diversity per genome using a custom R script. We are defining nucleotide diversity per genome as the number of sites with the reference allele divided by the total number of sites. Only sites that met our filtering criteria and contained no missing data across *P. pratensis* genotypes were included. Results were plotted with ggplot2 in R.

### Data availability

The genome assembly and annotation are available from the European Nucleotide Archive (ENA) under BioProject PRJEB51672. The raw Illumina sequence data for the *Poa* population panel is available from NCBI Sequence Read Archive (SRA) under BioProject ID PRJNA730042. The code for the entirety of assembly, annotation, and population genetic analyses is documented at https://github.com/phillipsar2/poa_genome.

## Results and Discussion

### Species identification and validation

Herbarium vouchers for the *Poa* reference genome and two of the population panel genotypes were identified as *P. pratensis* by their morphology (Supplement 1). The *Poa* reference genotype can be further classified as subspecies *angustifolia*, characterized by narrower and involute leaf blades, usually with strigose hairs on the adaxial surface of blades. The blades of *P. pratensis* subspecies *angustifolia* are firmer and tend to be more consistently glaucous. The intravaginal shoots are often disposed in fascicles of more than one shoot, the inflorescences are generally narrower, and the spikelets are smaller than other *P. pratensis* subspecies (Soreng and Barrie 1999; Soreng 2007; Cope and Gray 2009). *P. pratensis* subspecies *angustifolia* is the most likely classification for the reference genotype, although the infraspecies structure is complex and the subspecies genetically and morphologically grade into one another (Soreng and Barrie 1999; Soreng 2007; Cope and Gray 2009).

The remaining *Poa* population genotypes did not survive long enough for detailed morphological identification. We identified the remaining genotypes, and confirmed the morphological IDs, using phylogenetic inference with three commonly used loci (ETS, ITS, TLF). The reference genome was identified as *P. pratensis* by all three loci(Figures S1-3). Seven of the 8 genotypes in the *Poa* population panel were identified as *P. pratensis* by two of the three loci (ITS and ETS; Figures S1-2; Supplement 1) and held an unresolved position within the subgenus *Poa* in the third tree (TLF; Figure S3). The eighth population panel genotype was identified as *P. compressa* L. by all three loci. Phylogenetic identification thus supports our morphological identification of the reference genome as *P. pratensis*.

### Genome size and ploidy estimation

The reference individual was estimated to be octoploid given a genome size estimate of 3,525 Mbp and chromosome count of 54, assuming a basic chromosome number of *x* = 7 and a loss of two chromosomes (Figure S4; Supplement 5; Avdulov 1931; GPWG 2001). Further cytological studies are required to understand whether the chromosome loss is due to deletion or rearrangement. Our genome size estimate falls within the large range of genome sizes reported for *P. pratensis*, 2 to 9 pg/1C (Eaton *et al*. 2004; Huff and Bara 1993; Barcaccia *et al*. 1997; Raggi *et al*. 2015).

We also estimated the genome size of four of the eight population panel individuals. Genome size ranged from 3,248 to 4,856 Mbp with genotypes from the same population having similar genome sizes (Supplement 5). The substantial range in genome size variation in the population panel is not unexpected as *P. pratensis* is a polyploid series with common aneuploidy (Huff 2010). Given the range in the population panel, it is likely the genotypes have different chromosome counts and ploidy.

### Genome assembly

Error-corrected PacBio reads (100 Gb; 70X coverage) were assembled into 27,953 contigs. The contig assembly was oriented and further scaffolded using a Bionano optical map resulting in 118 primary scaffolds and 10 alternate scaffolds (Table 1).

**Table 1.** Assembly statistics.

| Variable | Description |
| --- | --- |
| Scaffolds | 118 |
| Contigs | 8,391 |
| Estimated genome size | 3.521 Gbp |
| Assembled genome size | 6.09 Gbp |
| Scaffold N50 | 65,127,037 bp |
| Scaffold L50 | 31 |
| Contig N50 | 1,095,498 bp |
| Contig L50 | 1548 |
| Longest scaffold | 177,118,352 bp |
| Scaffolds > 1 Mb | 110 |
| Scaffolds > 10 Mb | 98 |
| Average scaffold length | 51,622,171 bp |
| Average length of gaps | 44,233 bp |
| Complete BUSCOs | 99.2% |
| LAI | 25.8 |

The assembly is approximately 173% of the genome size (Table 1). Completeness of the assembly was assessed using Benchmarking Universal Single-Copy Orthologs (BUSCO) and the LTR Assembly Index (LAI). The assembly contains 99% of the expected conserved genes (BUSCOs), 98% of which were duplicated, and a LAI value of 25.8 indicates the transposable element assembly is also complete (Ou *et al*. 2018). Given the assembled genome size is approximately two-times the size of the estimated genome size and nearly all detected BUSCOs are duplicated, two unphased haplotypes are likely present in the assembly. Additionally, the high rate of duplicated BUSCOs may also be due to similarity among *Poa* subgenomes.

### Genome annotation

We identified 256,281 gene models, approximately 32K per subgenome assuming octoploidy, using a hybrid gene prediction pipeline that combined *ab initio* gene models with direct evidence annotations. Phylostrata demonstrated approximately 13% of the gene models are species-specific, which is higher than would be expected from orphan genes alone (Arendsee *et al*. 2014). Since the phylostratr program uses full proteomes from Uniprot to classify genes to their phylostrata, and there is lack of high-quality representative genomes for this clade, we observed an excess of species-specific genes. This demonstrates the important gap a *P. pratensis* reference genome fills in the green tree of life.

Transposable elements were comprehensively annotated using EDTA (Ou *et al*. 2019) and found to compose 58% of the genome. More specifically, Class I LTR retrotransposons and Class II DNA transposons comprise 36% and 15% of the genome, respectively. At the level of superfamily, the RLG (*Ty3*) LTR retrotransposon superfamily was the most common at 18% of the genome.

### Application of the reference genome

The reference genome contains multiple unphased haplotypes, and care should be taken in analyses that require genotypes or allele frequencies. Briefly, we discuss an alternative framework for estimating allele frequencies and potential pitfalls. Diploid genotypes (AA, Aa, aa) should not be called, as at least two haplotypes are assembled for many reference positions. Instead, we utilized an approach in which we randomly sampled a read from each position (Green *et al*. 2010). The randomly sampled read can then be used to calculate population allele frequencies and pairwise genetic distance matrices that are unbiased to sequencing depth or ploidy (Green *et al*. 2010; van der Valk *et al*. 2021; Pečnerová *et al*. 2021). Although we don’t detect a bias due to ploidy or chromosome count in our analyses (see below), these factors should always be considered in interpretation of results.

### Population genetics of North American Poa

Here, we demonstrate the effectiveness of the reference genome and a single-read genotyping approach in the estimation of population structure, using PCA and nucleotide diversity.

A PCA was run separately for all *Poa* genotypes, using 74,876 sites, and only *P. pratensis* genotypes, using 140,458 sites. The single-read genotypes were generated three times for the same set of sites and demonstrated similar results. We present the results for one run here. In the PCA with all *Poa* samples, most genetic variation was explained by species (27.9%) followed by population (16.2%; Figure 1A). *P. compressa* is distantly related to *P. pratensis* (Figure S1-S3) therefore we would expect the first principal component (PC) to separate by species. The second PC separates the *P. pratensis* genotypes in the Colorado population from two Manitoba *P. pratensis* genotypes (Figure 1A), while genotypes from the Colorado population remain clustered. The third principal component further separates the three *P. pratensis* populations.

**Figure 1.**
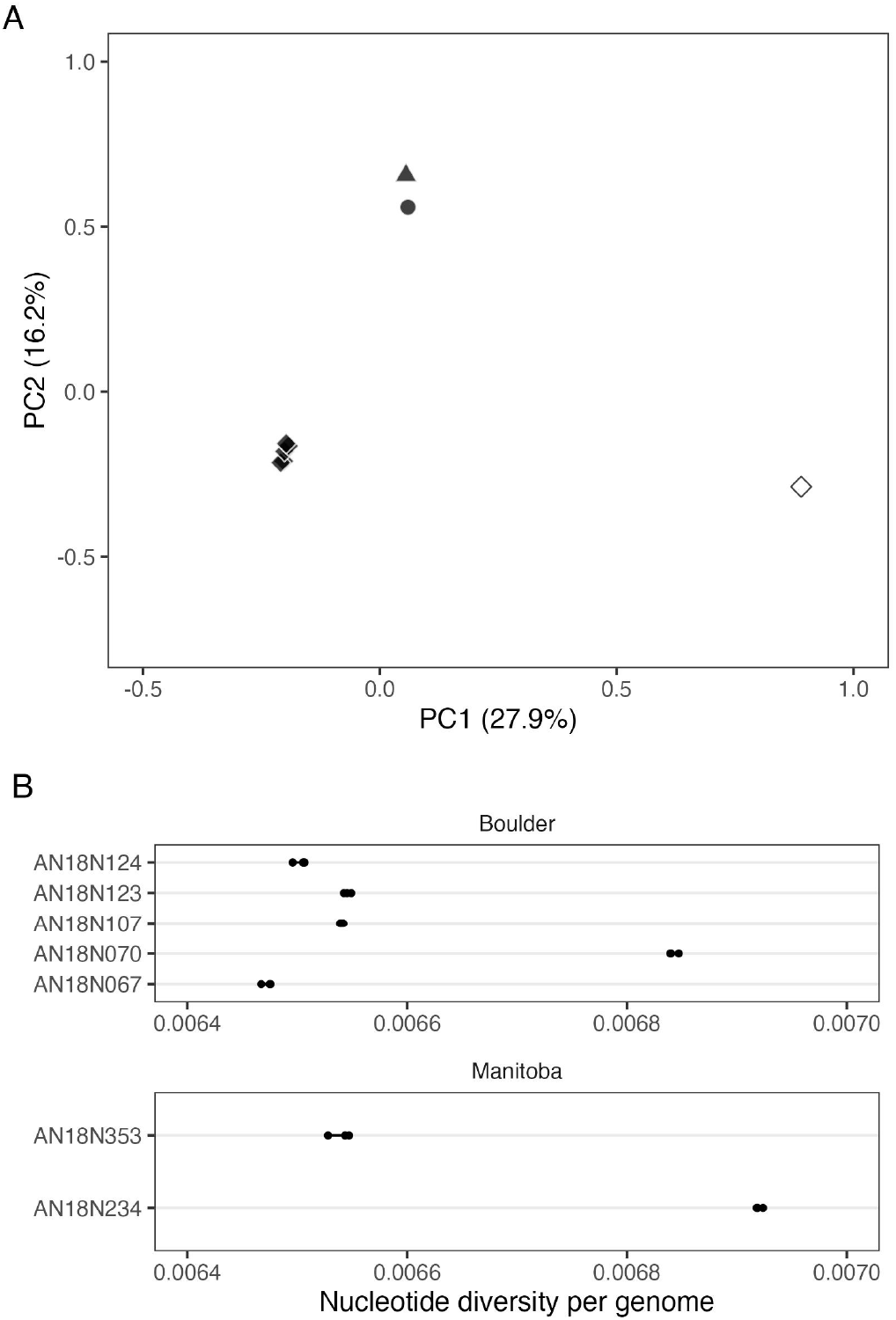
Population structure of *Poa* and nucleotide diversity in *P. pratensis*. (A) The first two PCs of a PCA of all sequenced *Poa* genotypes. The percent of genetic variation explained by each PC is reported in parenthesis on each axis. Sample locations are indicated by shape (circle = Argyle, Manitoba, triangle = Tolstoi, Manitoba, diamond = Boulder, Colorado) and species are colors (white = *P. compressa*, black = *P. pratensis*). (B) Mean nucleotide diversity per genome for only *P. pratensis* genotypes. Mean diversity of each run is plotted as a black circle for all genotypes.

The *P. pratensis*-only PCA demonstrates similar results with the first PC (24.6%) separating the Colorado genotypes from the two genotypes from Manitoba (Supplementary figure S5). The second PC (15.8%) separates the two genotypes from Manitoba and separates one Colorado genotype from the cluster. These results suggest North American *P. pratensis* populations are genetically differentiated and exhibit population structure, rather than being highly homogeneous or clonal. Our results support previous findings of population divergence in Northern Great Plains populations (Dennhardt *et al*. 2016).

To further understand the structure of genetic diversity across *P. pratensis* populations and the clustering within the Colorado population, we estimated nucleotide diversity per genome using 20,149,358 sites. Single-read genotypes were randomly drawn and nucleotide diversity was calculated three times with little variation between runs (Figure 1B; average variation between runs = 2.85*x*10^*−*11^). Mean diversity across *P. pratensis* genotypes is high (*π* = 0.0066, SD = 0.00017), which is consistent with previous studies of *P. pratensis* (Bonos and Huff 2013; Raggi *et al*. 2015; Bushman *et al*. 2013; Honig *et al*. 2018, 2012). The range of mean nucleotide diversity per genome within the Colorado population (0.0065 - 0.0068) and between the Manitoba genotypes (0.065 - 0.0069) is large, suggesting high within-population diversity.

## Conclusions

*Poa pratensis* is a globally popular turfgrass species used in lawns and recreation areas. Despite its economic value, progression of molecular tools to aid breeding has been slow compared to other turfgrasses as a result of polyploidy and apomixis (Bushman and Warnke 2013). Utilizing long read technology and a Bionano optical map, we have assembled and annotated the first high quality *P. pratensis* reference genome. We demonstrated the utility and application of the reference genome by evaluating the genetic diversity and population structure of wild North American *Poa*. As a result, we provided the first estimate of nucleotide diversity in *P. pratensis*.

Since our initial manuscript submission and preprint, Robbins *et al*. (2023) have published the genome of *P. annua*, a distantly related *Poa* species known as a weed and turfgrass worldwide. Future analyses, beyond the scope of this paper, comparing the two genomes will likely be fruitful for understanding the global success of *P. pratensis* and *P. annua*. As such, the *P. pratensis* reference genome and annotation will serve as an important resource in the study of bluegrasses.

## Acknowledgments

This project was supported by the National Science Foundation grant number 1822330. Thank you to Dr. Chrissy McAllister and Bess Bookout for sharing samples collected with permission from Nature Conservancy Canada properties and Lynn Riedel for collection of samples with permission from the City of Boulder Open Space and Mountain Parks. The Texas Advanced Computing Center (TACC) at The University of Texas at Austin, HPC@ISU equipment at Iowa State University (partially funded by NSF under MRI grant number 1726447) provided HPC resources that have contributed to the research results reported within this paper. We thank Dr. Kevin Fengler (for providing assembly instructions) and Dr. Gina Zastrow-Hayes (for establishing sequencing contracts), of Corteva Agriscience for their help in this project. A.R. Phillips would like to thank Andrew L. Murray for his support throughout the duration of this project. Additionally, the authors would like to thank our *Andropogon gerardi* reference plant for being contaminated with *Poa* and Felix Andrews for his alleged role in the happy accident that led to this work. Finally, thank you to Bob Ross for inspiring a generation of scientists to persevere.

## Supplement

**Figure S1.**
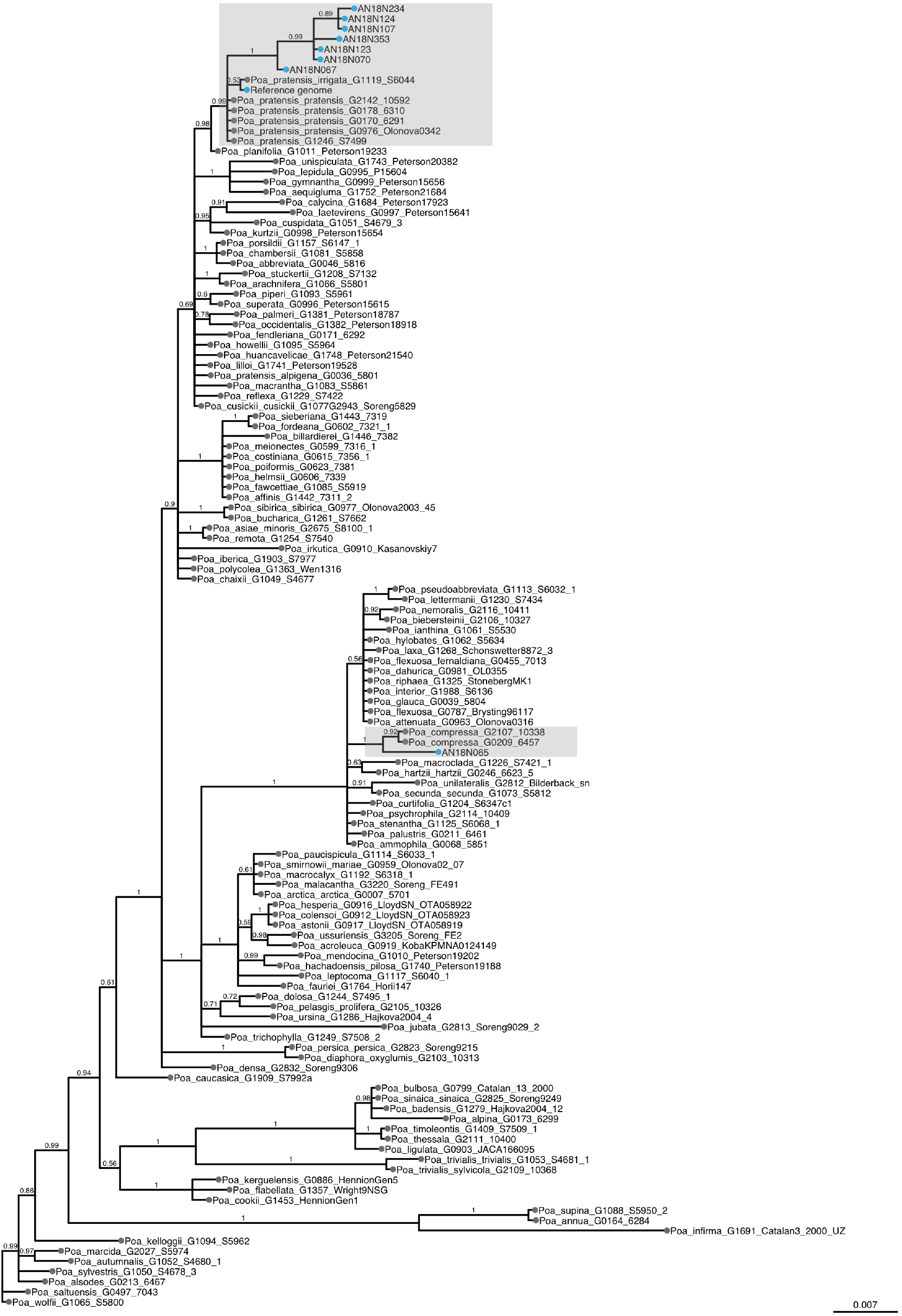
Bayesian 50% majority rule Consensus tree of ITS data. The *Poa* population panel and reference genome are indicated on the tree with blue dots. The unknown *Poa* population samples are labeled with their sample IDs (beginning with ‘AN’). The shaded boxes indicate the two clades the reference genome and population panel group within: *P. pratensis* and *P. compressa*. Bayesian posterior probabilities shown above the branches and branch length is the expected substitutions per site.

**Figure S2.**
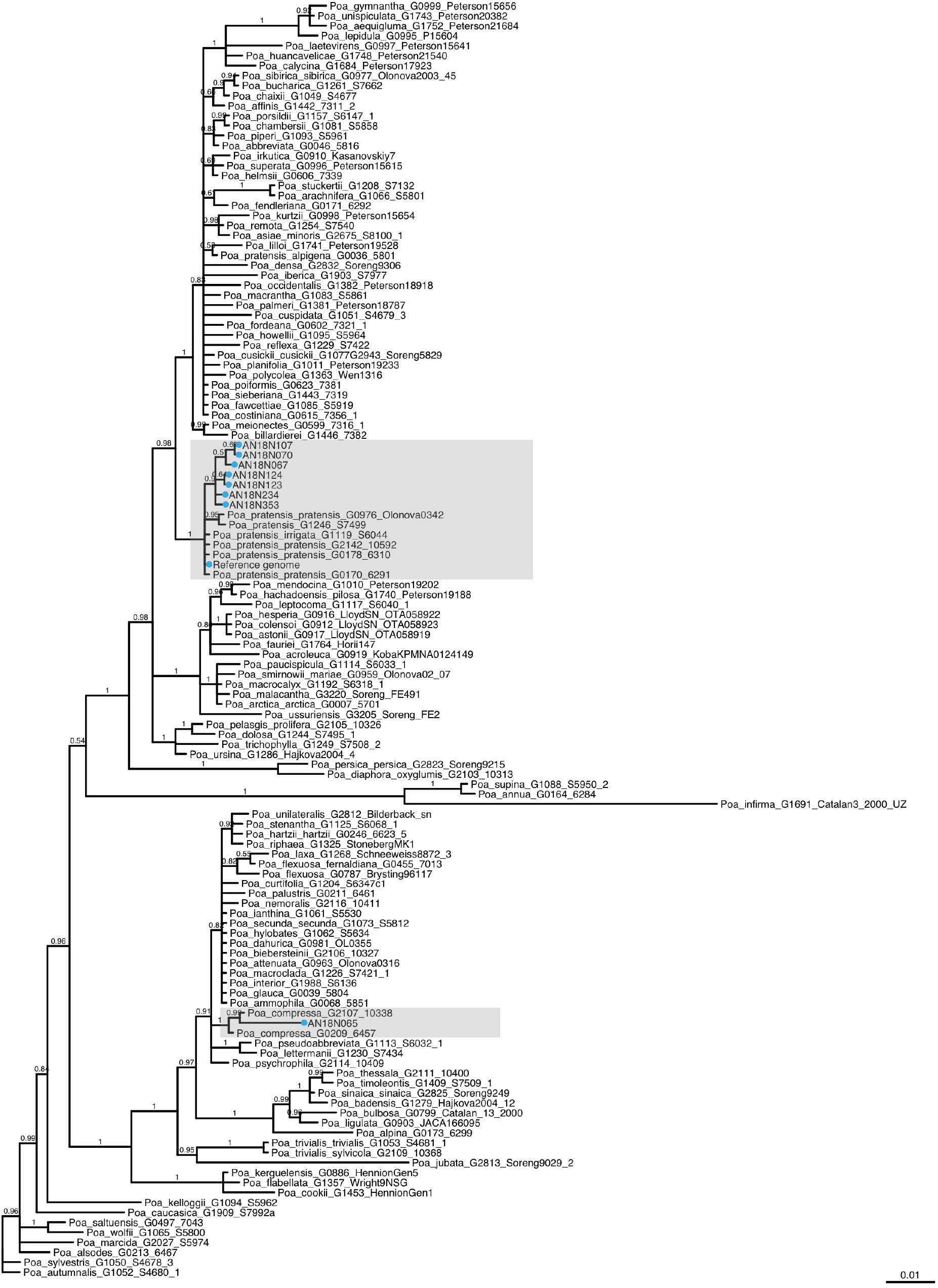
Bayesian 50% majority rule consensus tree of ETS data. See Supplement S1 for description of the figure components.

**Figure S3.**
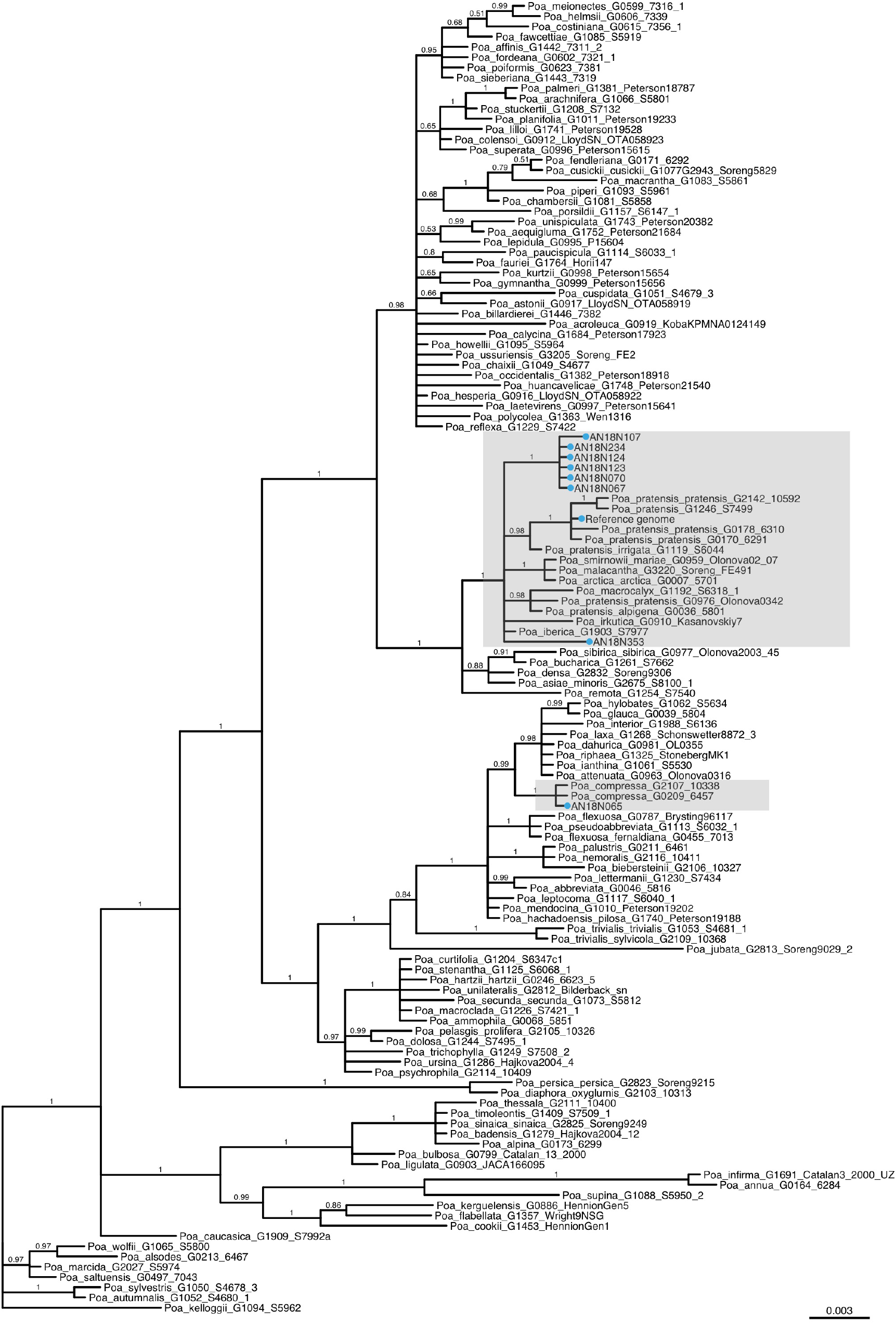
Bayesian 50% majority rule Consensus tree of TLF data. See Supplement S1 for description of the figure components.

**Figure S4.**
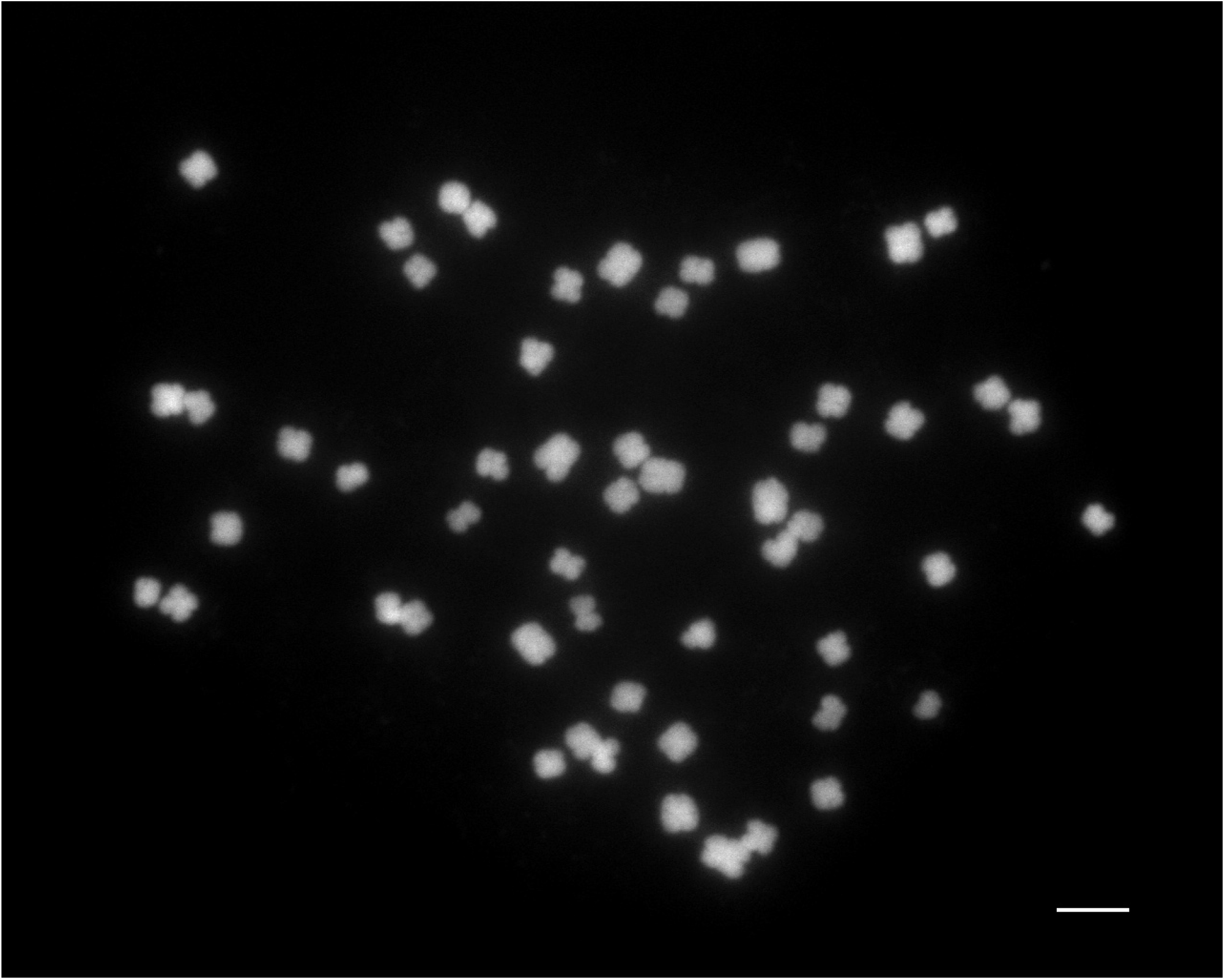
Metaphase chromosome spread with 54 chromosomes. *P. pratensis* reference individual root meristem cell counterstained with DAPI. Scale bar = 5 µm.

**Figure S5.**
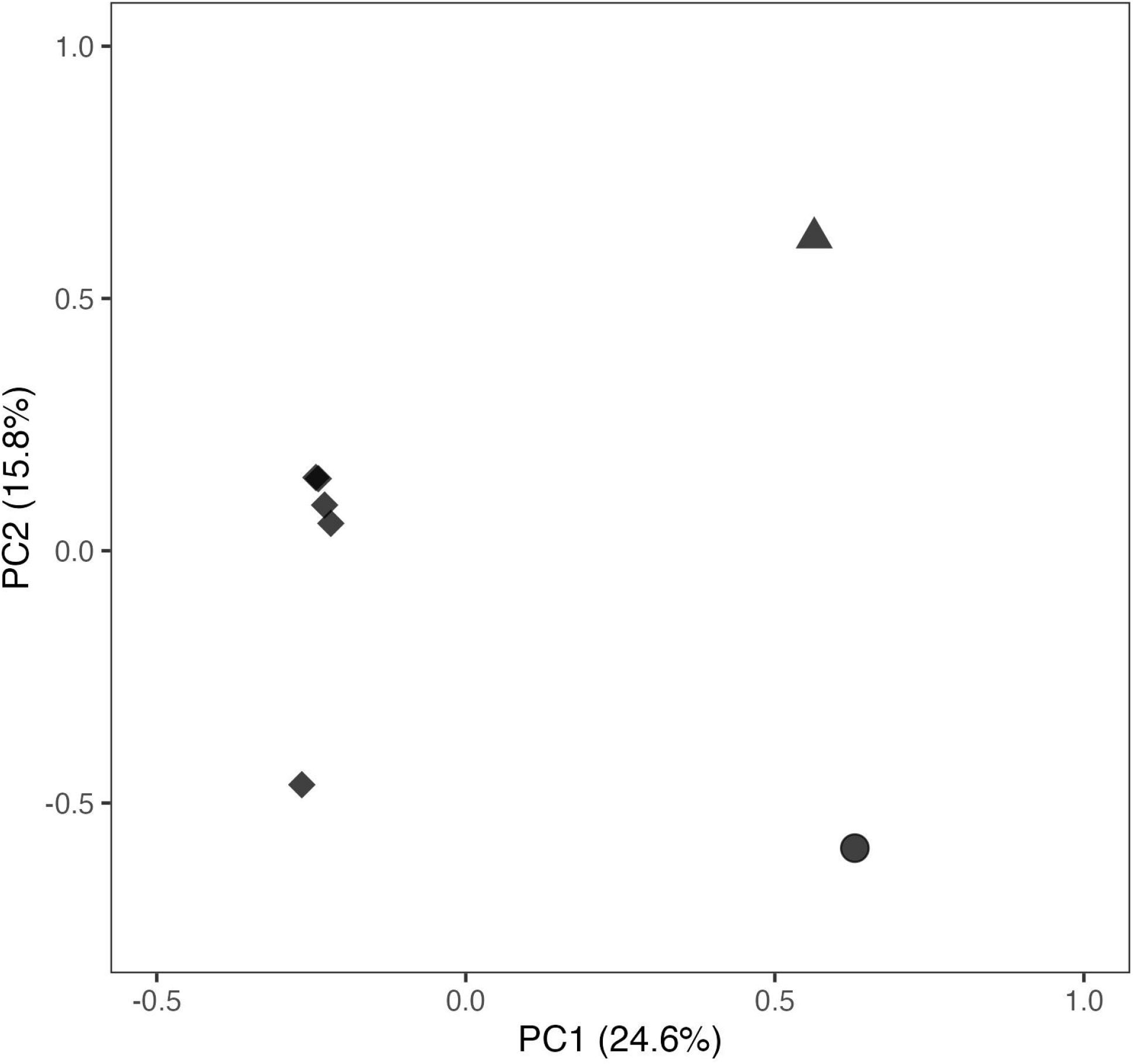
Population structure of *P. pratensis* genotypes only. The first two principal components (PCs) of a PCA of only the *P. pratensis* genotypes. The percent of genetic variation explained by each PC is reported in parenthesis on each axis. Sample locations are indicated by shape (circle = Argyle, Manitoba, triangle = Tolstoi, Manitoba, diamond = Boulder, Colorado).

